# Phylogenomic insights into distribution and adaptation of Bdellovibrionota in marine waters

**DOI:** 10.1101/2020.11.01.364414

**Authors:** Qing-Mei Li, Ying-Li Zhou, Zhan-Fei Wei, Yong Wang

## Abstract

Bdellovibrionota is composed of obligate predators that can consume some gram-negative bacteria inhabiting various environments. However, whether genomic traits influence their distribution and marine adaptation remains to be answered. In this study, we performed phylogenomics and comparative genomics studies on 82 Bdellovibrionota genomes along with five metagenome-assembled genomes (MAGs) from deep sea zones. Four phylogenetic groups, Oligoflexia, Bdello-group1, Bdello-group2 and Bacteriovoracia, were revealed by constructing a phylogenetic tree, of which 53.84% of Bdello-group2 and 48.94% of Bacteriovoracia were derived from ocean. Bacteriovoracia was more prevalent in deep sea zones, whereas Bdello-group2 was largely distributed in the epipelagic zone. Metabolic reconstruction indicated that genes involved in chemotaxis, flagellar (mobility), type II secretion system, ABC transporters and penicillin-binding protein were necessary for predatory lifestyle of Bdellovibrionota. Genes involved in glycerol metabolism, hydrogen peroxide (H_2_O_2_) degradation, cell wall recycling and peptide utilization were ubiquitously present in Bdellovibrionota genomes. Comparative genomics between marine and non-marine Bdellovibrionota demonstrated that betaine as an osmoprotectant is probably widely used by marine Bdellovibrionota, meanwhile, all the marine genomes have a number of related genes for adapting marine environment. The chitinase and chitin-binding protein encoding genes were identified for the first time in Oligoflexia, which implied that Oligoflexia may prey a wider spectrum of microbes. This study expanded our knowledge on adaption strategies of Bdellovibrionota inhabiting deep sea and their potential usage for biological control.

**Importance:** Bdellovibrionota can prey gram-negative bacteria proposed as biocontrol agent. Available Bdellovibrionota genomes showed that most are from marine environment. However, vertical distribution and adaption of Bdellovibrionota in deep sea has not been reported. Our study of Bdellovibrionota revealed four groups (Oligoflexia, Bdello-group1, Bdello-group2 and Bacteriovoracia) and their distribution pattern in oceans. We also identified the genes for different phases of predation and adaptation in deep-sea environment. Moreover, Oligoflexia genomes contain more genes for carbohydrates utilization and particularly those encoding chitin-binding protein and chitinase. Our analyses of Bdellovibrionota genomes may help understand their special lifestyle and deep-sea adaptation.

## Introduction

Bdellovibrionota, previously known as Bdellovibrio and like organisms (BALOs), were gram-negative bacterial predators and were isolated from various environments. Bdellovibrionota are small vibrio-shaped bacteria with a polar flagellum (1). In 1965, the first free-living BALO, *Bdellovibrio bacteriovorus*, was reported, which lead to a discovery that predation process of Bdellovibrionota includes two steps, 1) attachment and degradation of prey cell wall with glycanases and 2) digestion of prey cellular components with exoenzyme (2). Using *Escherichia coli* as a prey model, experimental results showed that Bdellovibrionota could penetrate the host cell and then transform into bdelloplast before being released from the host cell (3). In 1979, Bdellovibrionota was found to be an agent against pathogens (4), indicating the usage of Bdellovibrionota as a new approach of biocontrol that fights against bacterial infections, particularly when MDR bacteria are increasingly concerned globally (5). MDR bacterial infections have been successfully treated with assistance of bacterial predation by Bdellovibrionota (6). The prey spectrum of Bdellovibrionota is recently expanded to gram-positive pathogenic bacteria such as *Enterococcus* (7).

The current Bdellovibrionota phylum consists of several classes according to the Genome Taxonomy Database (GTDB, release89) (8), such as Bacteriovoracia, Bdellovibrionota, Oligoflexia and others without a taxonomic assignment. They are inhabitants that can be identified in various environments, such as soil, freshwater (river, lake and spring), ocean, wastewater treatment bioreactor and sewage (9–12). In different ecological niches, Bdellovibrionota could live with a prey-dependent or prey-independent lifestyle (11). This provides a clue of metabolic flexibility of Bdellovibrionota, as well as different predatory mechanisms when invading periplasmic space or attaching to the external surface of prey bacterial cell (1, 13). As bacterial predators distributed in a wide variety of niches with diversified predatory approaches, ecological roles of Bdellovibrionota were probably underestimated.

The ocean is probably the biggest reservoir of bacteria. So far the reported obligatory predator bacteria mainly include *Micavibrio* (a genus of α-proteobacteria) and five families belonging to δ-proteobacteria (14). Proteobacteria is a super phylum that constitutes gram negative bacteria and dominates the microbial communities in dark deep-sea zones (15). Experiments had exhibited that *Halobacteriovorax*, affiliated with one group of Bdellovibrionota, could predate deep-sea bacterium *Vibrio parahaemolyticus* (gram-negative bacterium) (16). In nutrient-poor deep sea, gram-negative bacteria may meet difficulties in synthesis biopolymers of key cellular structural components, such as peptidoglycan. Therefore Bdellovibrionota can probably prey on more gram-negative bacteria, which likely occurs ubiquitously in deep-sea environments (17). Under oligotrophic deep-sea environment, the competition for nutrient among microorganisms was constantly intensive (18). The adaptation strategy of Bdellovibrionota in marine environment is still an unresolved issue.

In this study, available Bdellovibrionota genomes were collected for phylogenomic analysis along with five representative Bdellovibrionota genomes binned from the marine metagenomes. The environmental sources of the genomes indicated that two subclades (Bacteriovoracia and Bdello-group2) were mostly present in marine environment. Bacteriovoracia were increasingly abundant in deeper ocean, while Bdello-group2 prefers to inhabit the epipelagic zone. Comparative genomics demonstrated that marine Bdellovibrionota species evolved to have genes involved in osmolyte metabolism. Oligoflexia may be a potential biocontrol agent of pathogens beyond bacteria since we discovered their chitin-binding protein and chitinase encoding genes.

## Results and discussion

### Phylogenomics of Bdellovibrionota

A total of 148 genomes of Bdellovibrionota were downloaded from GTDB database (Table S1), eight genomes were obtained from the NCBI to represent the model species that have been well studied previously (19, 20). After quality control (>70% completeness, < 10% contamination and average nucleotide identity ≽ 98.5%) (Tables S2 and S3), 82 genomes were retained for the subsequent study. Five MAGs that represent deep-sea Bdellovibrionota were retrieved from marine water samples of the Mariana Trench, Bashi Strait and the South China Sea with depths ranging from 1,646m to 5,992m (Table 1). The five deep-sea MAGs were in size of 2.19~5.88 Mbp with more than 70% completeness and less than 3% contamination (Table 1).

**Table 1.** Information of metagenome-assembled genomes of marine Bdellovibrionota.

| Bin_id | K1B34 | M27B12 | M27B96 | R3B4 | R4B59 |
| --- | --- | --- | --- | --- | --- |
| Depth (m) | 1,000 | 3,000~5,000 | 3,000~5,000 | 1,646 | 5,992 |
| Sampling site<br>(E; N) | 116.47°;<br>18.01° | 142.01°;<br>11.16° | 142.01°;<br>11.16 ° | 120.03°;<br>21.87° | 123.31°;<br>22.82° |
| Com. (%) | 75.16 | 98.21 | 80.15 | 81.83 | 71.94 |
| Con. (%) | 0.00 | 0.89 | 2.98 | 2.52 | 0.89 |
| N50 (bp) | 6,396 | 295,316 | 5,991 | 7,831 | 3,968 |
| No. contigs | 597 | 24 | 684 | 923 | 570 |
| Genome size<br>(bp) | 3,229,602 | 4,849,992 | 3,554,382 | 5,883,831 | 2,188,444 |
| No. CDSs | 3,741 | 4,431 | 4,172 | 5,667 | 2,566 |

\* R and K in Bin\_id represent the South China Sea; M denotes the Mariana Trench.
Com., completeness; Con., contamination;
N50, the shortest contig length required to cover 50% of the genome. CDS, coding
sequence.

Construction of phylogenomic tree using conserved proteins displayed four phylogenetic groups (subclades), which were then named as ‘Bacteriovoracia’, ‘Oligoflexia’, ‘Bdello-group1’ and ‘Bdello-group2’ (Figs. 1A and 1B). The phylogenetic relationships between the groups were consistent with those based on 16S rRNA genes (Fig. S1). The five deep-sea MAGs were distributed into three Bdellovibrionota groups except for Bdello-group1 (Figs. 1A and 1B). The available isolation sources of the samples and genomes were collected and summarized (Table S4). These genomes were mainly obtained from marine water, subsurface sediment and bioreactor sludge, waste water and ground water. Bdello-group2 and Oligoflexia were found in more diversified environments (Fig. 1C). Marine and ground water were the most common sources for Bdellovibrionota, which coincides with their preference for environment with low viscosity (21). About 43.40% of Bacteriovoracia and 46.81% of Bdello-group2 genomes were derived from marine waters (Fig. 1C and Table S4), implying their important role in oceans. The wide spread of Bdellovibrionota may be attributable to two-phase lifestyle and quick response to the transformation between the phases (22).

**Figure 1.**
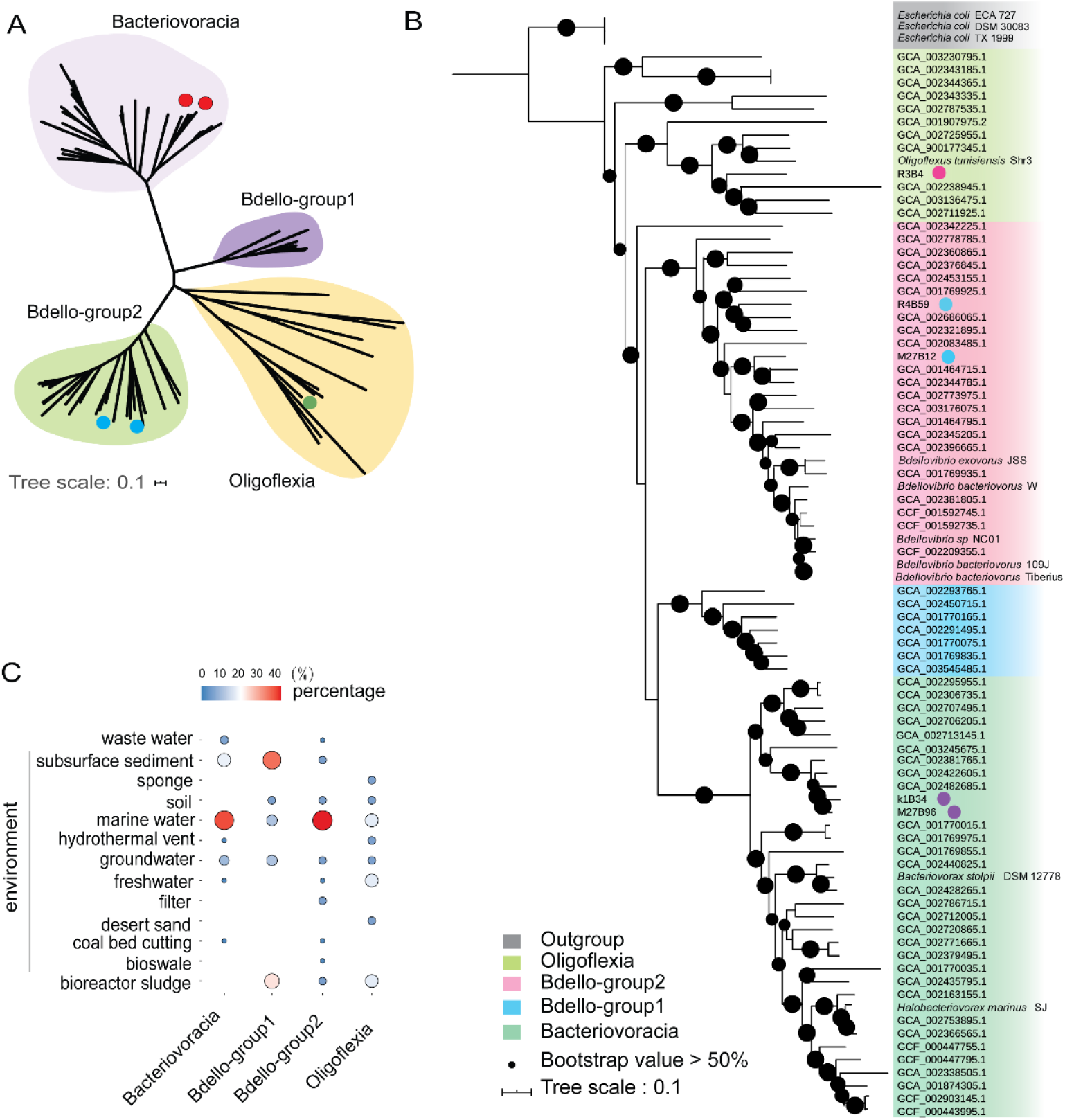
Phylogenomic analysis and distribution of Bdellovibrionota. (A) Unrooted maximum-likelihood (ML) phylogenomic tree of Bdellovibrionota. The ML tree was constructed based on 43 concatenated conserved proteins with the PROTGAMMALG model. The MAGs from this study were marked with a dot. (B) Rooted ML phylogenomic tree of Bdellovibrionota. The tree was built up using three *E. coli* genomes as a outgroup. The genomes with a dot were marine MAGs from this study and the references were shown as their GTDB accession numbers or species names. (C) Environmental source of Bdellovibrionota genomes. The size and color of dots represent the percentage of the sources in each group.

Among the four groups, the size of Oligoflexia genomes varies considerably (Fig. 2A). This is true for their genomic GC contents ranging from 32.83 % to 54.35% (Fig. 2A and Table S2), which might be stemmed from the presence of many copies of transposase genes in their genomes (Fig. S2). The average GC content of Bdello-group1 genomes was highest with a mean value more than 50% (Fig. 2A and Table S2). Note that 26.67% of the Bdello-group1 genomes were obtained from sludge bioreactors (Table S4). Whether the wide range of GC contents of Bdello-group1 genomes is ascribed to frequent contacts with the diverse microbes in sludges is still elusive.

**Figure 2.**
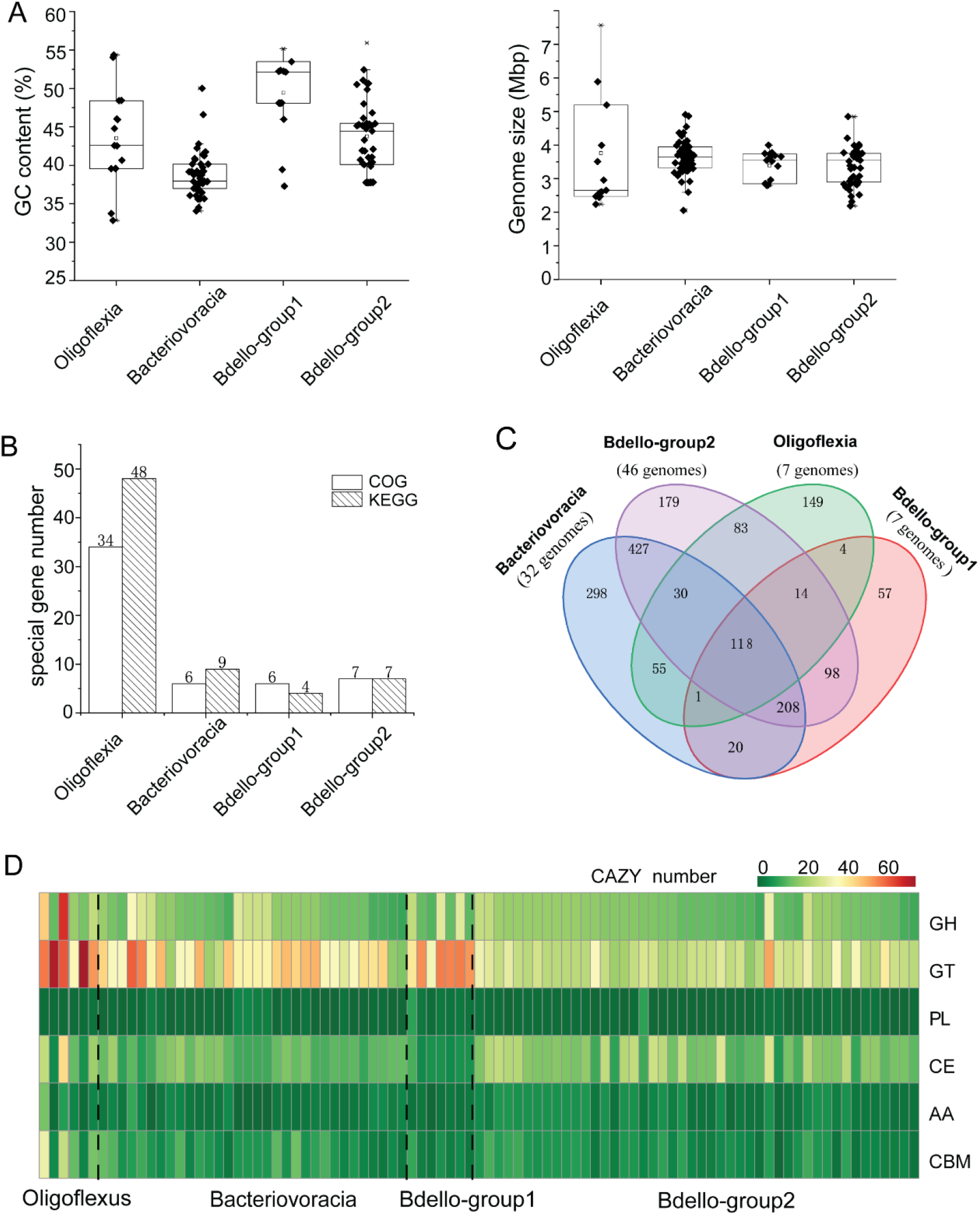
Genomic and gene content difference among Bdellovibrionota groups. (A) GC content and genome size of Bdellovibrionota genomes. (B) Number of group-specific COGs and KEGG genes. (C) Venn diagram showing group-specific and shared peptidases of Bdellovibrionota. (D) Heatmap illustrating differences in number of carbohydrate active enzyme (CAZy) classes.

### Relative abundance of Bdellovibrionota in marine water zones

To examine vertical distribution of the four Bdellovibrionota groups (Fig. 1A) in marine water zones, the 16S rRNA sequences (16S metagenomic Illumina tags, miTags) of Tara ocean (23) and Mariana marine metagenomes (24, 25) were used to calculate relative abundance of Bdellovibrionota groups. Bacteriovoracia and Bdello-group2 were relatively abundant in oceans, compared to the other two groups (Fig. 3A). The relative abundance of Bacteriovoracia increased with water depth, whereas a reverse trend was observed for Bdello-group2. Statistical analysis showed that epipelagic layer had significantly more abundant Bdello-group2 than the other deep-sea layers, whereas significantly more abundant Bacteriovoracia were distributed in deeper layers (t-test, *p*<0.05) (Fig. 3B). These results strongly indicate that Bacteriovoracia was relatively more prevalent in deeper marine environment. Conversely, Bdello-group2 prefers to inhabit the epipelagic zone. Oligoflexia and Bdello-group1 are associated with very low relative abundance in marine environment (Fig. 3B). Oligoflexia showed increased tendency with significance between epipelagic and mesopelagic zones (t-test, *p*<0.05), whereas there was no significance when comparing the epipelagic and deeper layer (≽ 1,000m) (Fig. 3B). Those results showed that vertical distribution pattern of Bdellovibrionota differed between the groups, suggesting considerable difference in their gene profiles.

**Figure 3.**
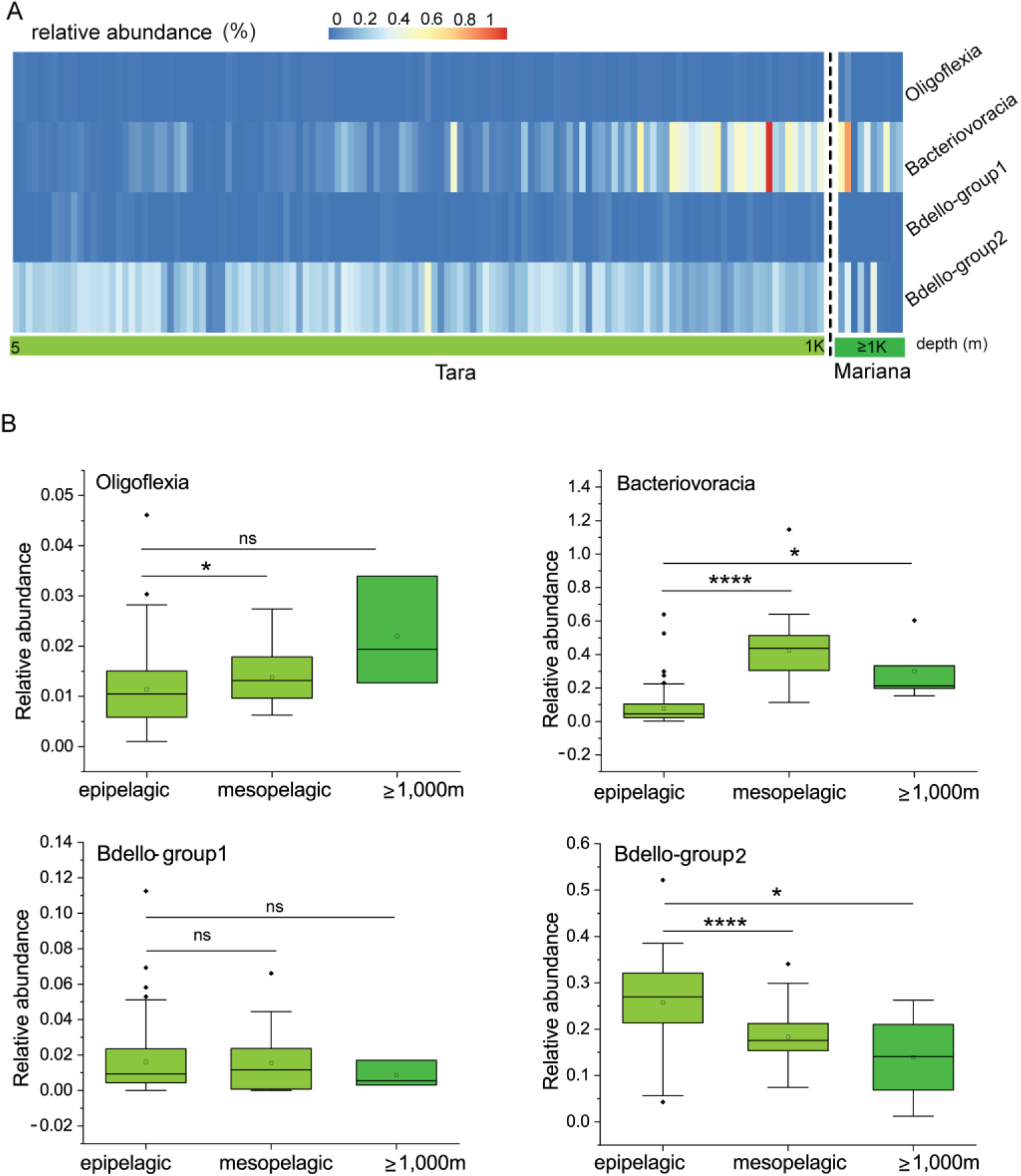
Vertical distribution of Bdellovibrionota in marine waters. (A) Relative abundance of Bdellovibrionota was calculated as percentage of Bdellovibrionota miTags in metagenomes of the Tara Ocean project (depth from 5m to 1,000m) and the Mariana Trench (≽ 1,000m) (Table S7) (Gao *et al.*, 2019; Li *et al.*, 2019). (B) Relative abundances of Bdellovibrionota groups in different marine layers were plotted. T-test was performed to compare the relative abundances at different layers. (****,*p*<0.0001; *,*p*<0.05).

### Metabolic potentials (reconstruction) of Bdellovibrionota

Discrepancies of the Bdellovibrionota groups in metabolism and environmental adaptation were detected by comparing annotations against KEGG (Kyoto Encyclopedia of Genes and Genomes), COG (Cluster of Orthologous Groups of proteins), MEROPS and CAZY (Carbohydrate Active enZYmes) databases (26–29). When group-specific genes (only present in more than 50% genomes of one group) were selected from KEGG and COG annotation results, we found that Oligoflexia genomes contained 48 KEGG genes and 34 COGs as group-specific genes, remarkably more than other groups (Fig. 3B). The peptidase-coding genes found in MEROPS annotation were compared, which showed that 118 peptidase genes were shared by four groups and each group also contained unique peptidases (Fig. 3C). Oligoflexia differs from other groups of Bdellovibrionota with higher average number of unique peptidases (Fig. 3C). Based on CAZY annotation, Oligoflexia genomes encode the obviously more enzymes of glycosyl transferase (GT) families than other groups (Fig. 3D). GT enzymes can catalyze the transfer of sugar moieties onto aglycons, which is related to transformation of many natural products (30). This indicates that Oligoflexia may heavily depend on degradation of carbohydrate from prey bacterial cells and environment.

To disentangle the metabolic and adaptive capacities of Bdellovibrionota, metabolism reconstruction was carried out and presented (Fig. 4). If at least 50% genomes of a group harbor a gene, the group was regarded to have the gene. As bacterial predators, Bdellovibrionota genomes all harbor genes that encode an almost full set of subunits involved in mobility (FlgKGI, FliL and MotA), chemotaxis (CheYVBWAR and MCP), type II secretion system and some ABC transporters (Table S5). These chemotactic proteins may work effectively and help Bdellovibrionota to sense abundance of prey bacteria and toxic chemicals. Multicopies of *mcp* and *motA* genes were predicted in the genomes (up to 27 copies of *mcp* in an Oligoflexia genome, GCA_001907975.2, renamed as ref32 in this study) (Table S6). Artificial mutations of *motA* affected integrality of flagellar (31), which indicated that MotA is important in flagellar assembly. MCP, a methyl accepting chemotaxis protein, was reported to act as an enhancer for predatory ability of Bdellovibrionota (32). In addition, *cheB* was predicted in all the groups (71.42% of Oligoflexia genomes; all of the Bacteriovoracia genomes; 57.14% of Bdello-group1 genomes and 89.13% of Bdello-group2 genomes) (Table S5). During bacterial rapid adaption and responding to the local gradients of attractants or repellents, protein CheB, a methylesterase, functions in covalent modification of membrane receptors (33). A gene encoding a methyltransferase CheR is present in all the Bdellovibrionota groups (Table S5) (34). Methylation enhances clockwise signals of the motors, while demethylation attenuates them by means of a feedback loop that is managed by CheB and CheR with respect to methylation rate of receptors (35). This is the nature of motor transition between clockwise and counterclockwise to response the changes of environmental factors. Bdellovibrionota may hunt other bacteria with assistance of methylated modification related chemotaxis proteins (CheB, CheR and MCP) for precise control of bacterial motility.

**Figure 4.**
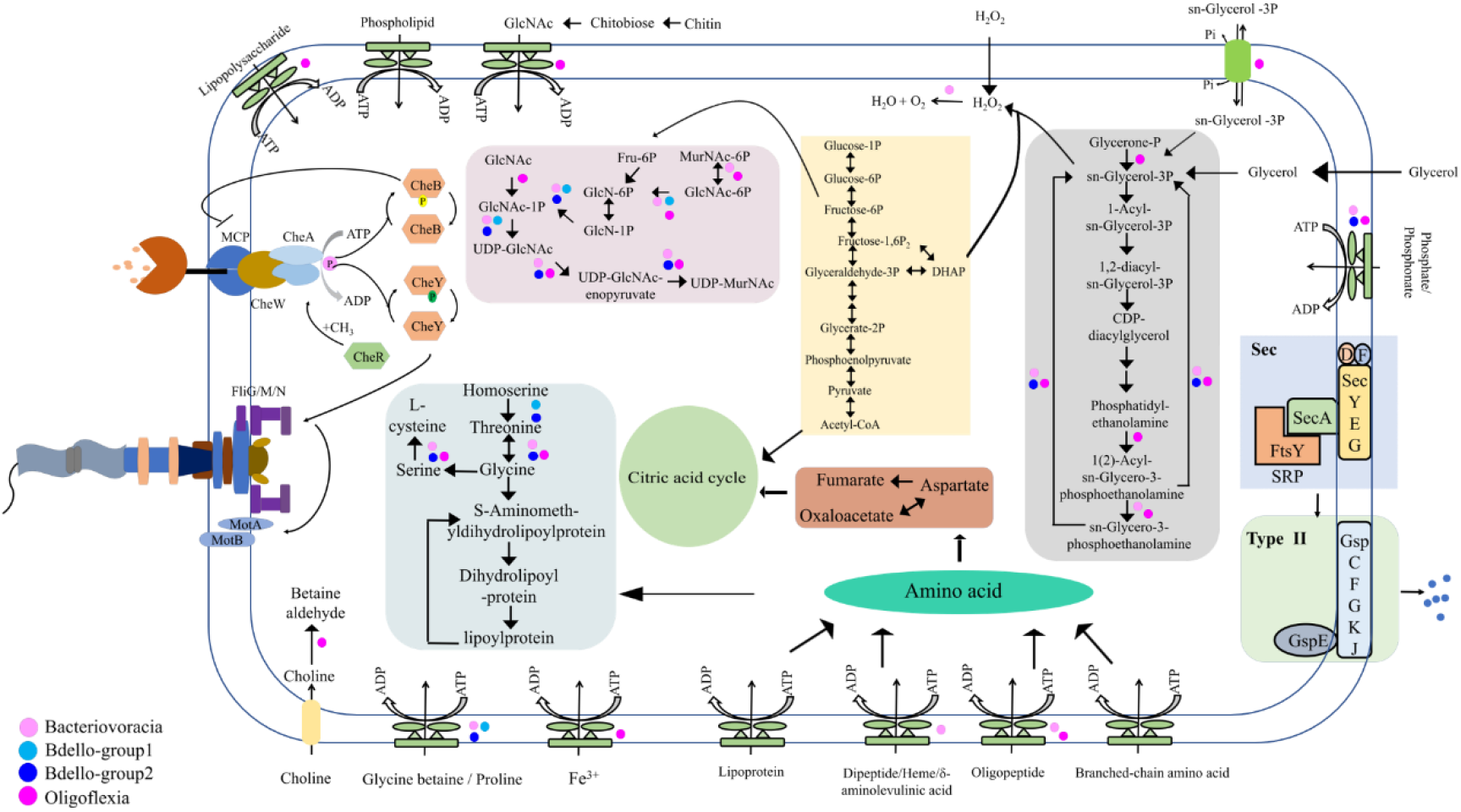
Metabolism reconstruction of Bdellovibrionota. The schematic metabolism and structural components of Bdellovibrionota were predicted based on KEGG annotation results for the four groups. The genes responsible for the steps without a dot are idenfitied in all the four groups of Bdellovibrionota; if not, dots were used to denote their presence in group(s).

After invasion into periplasmatic space of prey bacteria, Bdellovibrionota bacteria begin to degrade outer cellular structures and cell membrane of the prey. N-acetyl-D-glucosamine (glcNAc), glycerol and lipid of cellular components are all nutrients for Bdellovibrionota. After glcNAc is imported into a Bdellovibrionota cell, it can be directly used for producing UDP-N-acetylmuramic acid (UDP-MurNAc) for subsequent biosynthesis of peptidoglycan that is the basic unit of cell wall (Fig. 4) (36). GlcNAc may also be degraded into glycolysis pathway. Glycerol, a simple organic molecular, can be produced by several metabolism pathways (such as glycolysis and lipid catabolism). The coding genes of glycerol kinase (GK) and glycerol-3-phospahte-dehydrogenase (GLPD) were predicted in almost all of the Bdellovibrionota genomes (Table S6), which indicates that Bdellovibrionota can utilize glycerol for glycolysis (Fig. 4). The by-production of glycerol oxidation is H_2_O_2_, which is cytotoxic (37) and can be used to destroy cell wall structure of prey bacteria after free diffusion of H_2_O_2_ into the prey periplasmic space. The coding gene of sn-glycerol-3-phosphate acyltransferase (PlsB), the first enzyme for membrane phospholipid biosynthesis (38), was predicted only in Oligoflexia genomes (Table S6). This suggests that glycerol may be used for membrane phospholipid biosynthesis in rapid growth phase of Oligoflexia and thus the glycerol metabolism was diversified in Bdellovibrionota.

The metabolism reconstruction shows that there are numerous kinds of ABC transporters for uptake of phospholipid, lipoprotein, oligopeptide and branched-chain amino acids (Fig. 4). They are also useful to import nutrients from environment in free-living phase. The strategies may help Bdellovibrionota to adapt to oligotrophic, low temperature and hydrostatic pressure in deep sea zones.

The degradation and assimilation of nutrients are essential processes during predatory growth phase of Bdellovibrionota. When Bdellovibrionota invaded the prey bacteria and entered into growth phase, the transcription of genes increased from 33% to 85% (22). In Bdellovibrionota, amino acids derived from prey bacterial cell, environmental source and oligopeptide digestion were fed into different metabolism pathways. For example, aspartate is linked with citric acid cycle and glycine may involve into homoserine metabolism (Fig. 4). We then examined whether homoserine might be transformed to acyl-homoserine lactone as the substance for quorum sensing that can further trigger type II secretion system by searching relevant genes for downstream transformation (39). However, there are no predicted acyl-HSL synthases in Bdellovibrionota genomes (LuxR/LuxI, AinR/AinS and other homologs to acyl-HSL synthases) (Table S6), whereas the enzymes that catalyze homoserine to threonine, glycine, serine and L-cysteine were identified (Fig. 4). These results indicate that Bdellovibrionota may just use homoserine as an intermediate metabolite for amino acid metabolism. Because of the special lifestyle of Bdellovibrionota, there may be other protein factors serving as the quorum sensing regulators during the predatory growth phase. The latest study indicated that diffusible signaling factor (DSF) considerably delayed predation and bdelloplast lysis (40). Sec-type II system is almost complete in all Bdellovibrionota groups (Fig. 4 and Table S5). Type II secretion system can exude numerous bacterial toxic proteins and lytic enzymes such as proteases and lipases, which is important for bacteria to obtain nutrient from prey cell membrane and environment (41). Bdellovibrionota may heavily depend on their type II secretion system in the whole lifecycle. During attack phase, the secreting proteins can probably be used to catch prey bacteria and partly destruct the outer membrane structure for entry; during the predatory growth phase, they are likely useful for cellular component digestion and complete prey cell lysis.

### Genes involved in survival of Bdellovibrionota in marine environment

Although many Bdellovibrionota genomes were obtained from marine environment, there is not a special group exclusively consisting of marine species (Fig. 1). Given the characteristics with high hydrostatic pressure, oligotrophy and low temperature in deep-sea, there must be some genes in Bdellovibrionota for deep-sea adaptation. To reveal these genes, we divided each group of the Bdellovibrionota genomes into two parts, non-marine and marine, according to their sample sources. The marine genomes of Bdellovibrionota mostly belonged to Bacteriovoracia and Bdello-group2 (Fig. 1C and Table S4). A comparison between the two groups showed that the marine genomes have *betT/betS* (77.78% of Bacteriovoracia marine genomes and 57.89% of Bdello-group2 marine genomes) that takes part in betaine biosynthesis via choline pathway because of its high-affinity to choline (Fig. 5). Betaine is one of known major organic molecules that protect bacteria from damage of hypersaline/high hydrostatic pressure by stabilizing the natural conformation of proteins and preventing their aggregation (42). Although it is reported that choline and glycine betaine are nearly ubiquitous in bacteria (42), the percentage of *betT*/*betS* in the marine group is much higher than that in non-marine group (77.78% vs. 21.05% of Bacteriovoracia and 57.89% vs. 0 of Bdello-group2) in this study (Fig. 5). This indicates that *betT*/*betS* genes in the marine Bdellovibrionota may be critical in protecting them from the high hydrostatic pressure by importing choline/glycine/proline betaine from the sea water or the prey bacterial cells. Apart from *betT*/*betS*, the gene named *mscS* (Fig. 5), contributing to normal resistance to hypoosmotic shock, was predicted in 89.47% of Bdello-group2 marine genomes. The MscS channel is integrated in the cell membrane, which prevents burst of the cell in hypotonic environments (43, 44). It may work with BetT/BetS in a feedback model for maintaining and rescuing Bdellovibrionota (Bdello-group2) under the various marine chemical gradients.

**Figure 5.**
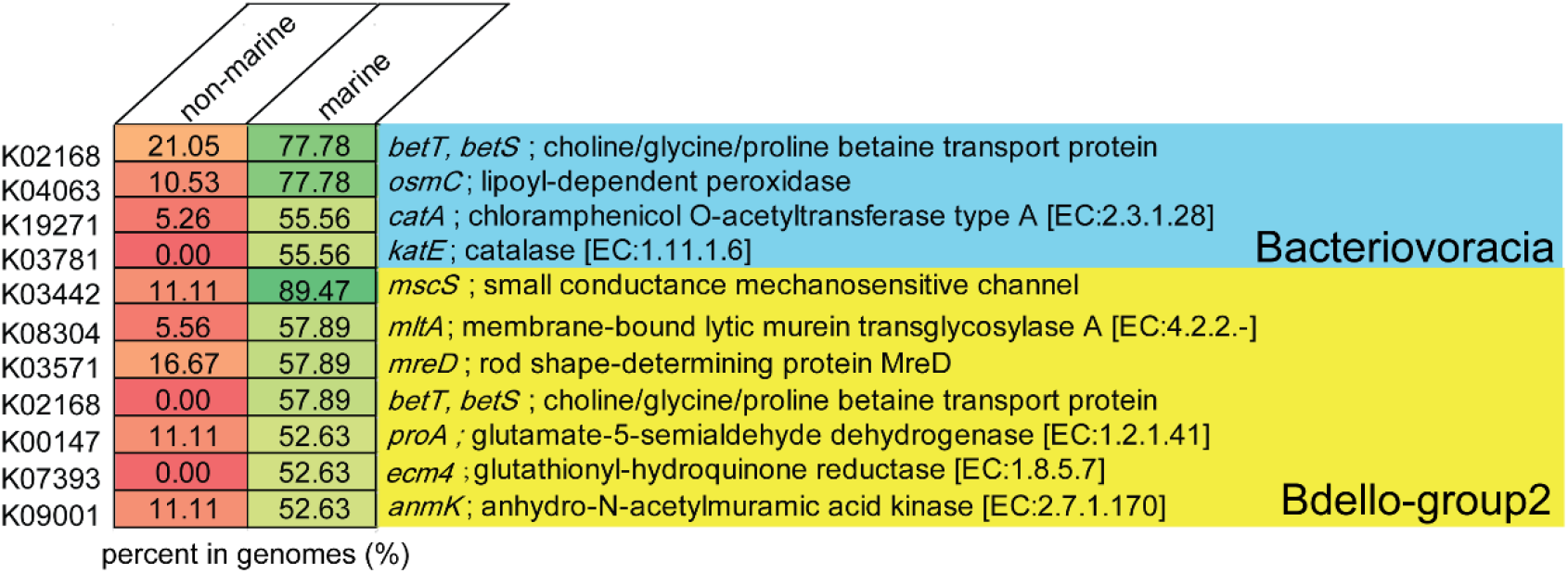
Genes enriched in marine genomes of Bacteriovoracia and Bdello-group2. The percentage of the KEGG genes in the genomes was calculated for the marine and non-marine genomes, respectively, in the two Bdellovibrionota groups.

The special physicochemical parameters of ocean lead to diverse resistance mechanisms of marine microbes, such as detoxification. The H_2_O_2_ degrading genes were predicted in Bacteriovoracia marine genomes, and were perhaps involved in prevention against self-toxification of H_2_O_2_ induced by the byproduct of glycerol oxidation. The genes *osmC* (77.78%), *catA* (55.56%) and *katE* (55.56%) were present in Bacteriovoracia marine genomes (Fig. 5). These genes are probably related to protect cells from the toxic effects of H_2_O_2_ that acts as a bactericide (45). OsmC is a protein induced by osmotic pressure (46). Interestingly, its malfunction reduced fitness and elevated sensitivity to oxidative stress of *E. coli* (47). Therefore, OsmC may participate in defense against high salinity and H_2_O_2_ (a defensive molecule that may also be produced by prey bacteria) in Bacteriovoracia (Fig.4). As for *catA*, it was up-regulated in *Streptomyces coelicolor* treated with H_2_O_2_ during no-stationary phase and it was required for *Streptomyces coelicolor* growth under aerobic condition (48, 49). The function of *katE* has not yet been determined, but it is predicted to regulate the concentration of H_2_O_2_ (50, 51). H_2_O_2_ may be scavenged and degraded in Bacteriovoracia with co-operation of OsmC, CatA and KatE for survival in deeper ocean (Figs. 3 and 5).

In Bdello-group2 marine genomes, there are some genes involved in cell wall degradation, recycling, organization and biogenesis (Fig. 5). Penicillin-binding proteins are important for Bdellovibrionota to break up prey cell wall (52). MltA encoded by 57.89% of Bdello-group2 marine genomes is a murein degrading enzyme that is located in outer-membrane and can interact with penicillin-binding protein 2 (53, 54), which is important for marine Bdello-group2 as the predator of other gram-negative bacteria. *MreD* gene was present in 57.89% of Bdello-group2 marine genomes (Fig. 5) and is essential for lateral peptidoglycan synthesis (55). We identified *anmK* gene in 52.63% of Bdello-group2 marine genomes (Fig. 5). The enzyme encoded by *anmK* plays a role in cell wall recycling though transforming 1,6-anhydro-N-acetylmuramic acid (anhMurNAc) to N-acetylglucosamine-phosphate (GlcNAc-P) (56). Usually, the source of anhMurNAc is derived from its own cell wall murein, but as predators, marine Bdello-group2 may reuse the component of prey bacteria for cell wall formation of next generation. ECM4 found in 52.63% of Bdello-group2 marine genomes (Fig. 5) was proposed as a protein involved in cell wall biogenesis (integrity) (57). Under deep-sea oligotrophic condition, these genes may be all associated with the whole process from murein-degrading, anhMurNAc recycling and cell wall biogenesis for proliferation of Bdello-group2 marine bacteria.

There are two genomes of Bdello-group1 (GCA_002722705.1 and GCA_002450715.1) and only one genome of Oligoflexia (R3B4) from marine. The discrepancies in genes between marine and non-marine groups were also demonstrated (Fig. S2). The genes involved in synthesis of osmoprotectants were also predicted in marine Bdello-group1 and Oligoflexia genomes, such as *betT* and *mscS* in Bdello-group1 and *cmo* in Oligoflexia (Fig. S2). CMO can catalyze the first step of glycine betaine synthesis (58), which indicates that glycine betaine can be used as an osmoprotectant in marine Oligoflexia. In addition, there are many genes involved in transport or transformation of sugar as osmoprotectant, such as *malK*, *msmX*, *rgpF* and *algI* genes in Oligoflexia (Fig. S2) (59–62). Combining these results, betaine as an osmoprotectant is commonly used among different groups of Bdellovibrionota, all having a variety of genes for fitness in ocean.

### Oligoflexia as potential defender against eukaryotic pathogens

In light of the comparison among Bdellovibrionota groups, Oligoflexia seems to be special in gene content. The chitinase gene (K01183) has many copies in Oligoflexia genomes (Fig. S3), but none in other Bdellovibrionota groups. The presence of genes encoding chitinase and chitin-binding protein (Fig. S3) indicates that some Oligoflexia may be able to degrade chitin (63, 64). The phylogenetic tree of the seven chitinases extracted from the Oligoflexia genomes and other species supports their close relationships with known homologs (Fig. 6). To further verify the predicted open reading frame (ORF) of the chitinases, multiple alignment was performed to validate the conserved motif ‘DXXDXDXE’ of chitinases (Fig. 6) (65). In addition, the Oligoflexia genomes contained a gene coding for chitin-binding protein (Fig. S4 and Table S8). These results imply that a part of Oligoflexia can utilize chitin in environment as nutrient and/or attack preys with chitin ultracellular structure. In this study, the chitinase and chitin-binding protein coding genes were focused for the first time in Oligoflexia. Pathogenic fungi have chitin as a main cell wall component (66), which allows to speculate that Oligoflexia may be used to defend against eukaryotic pathogens aside from bacterial pathogens.

**Figure 6.**
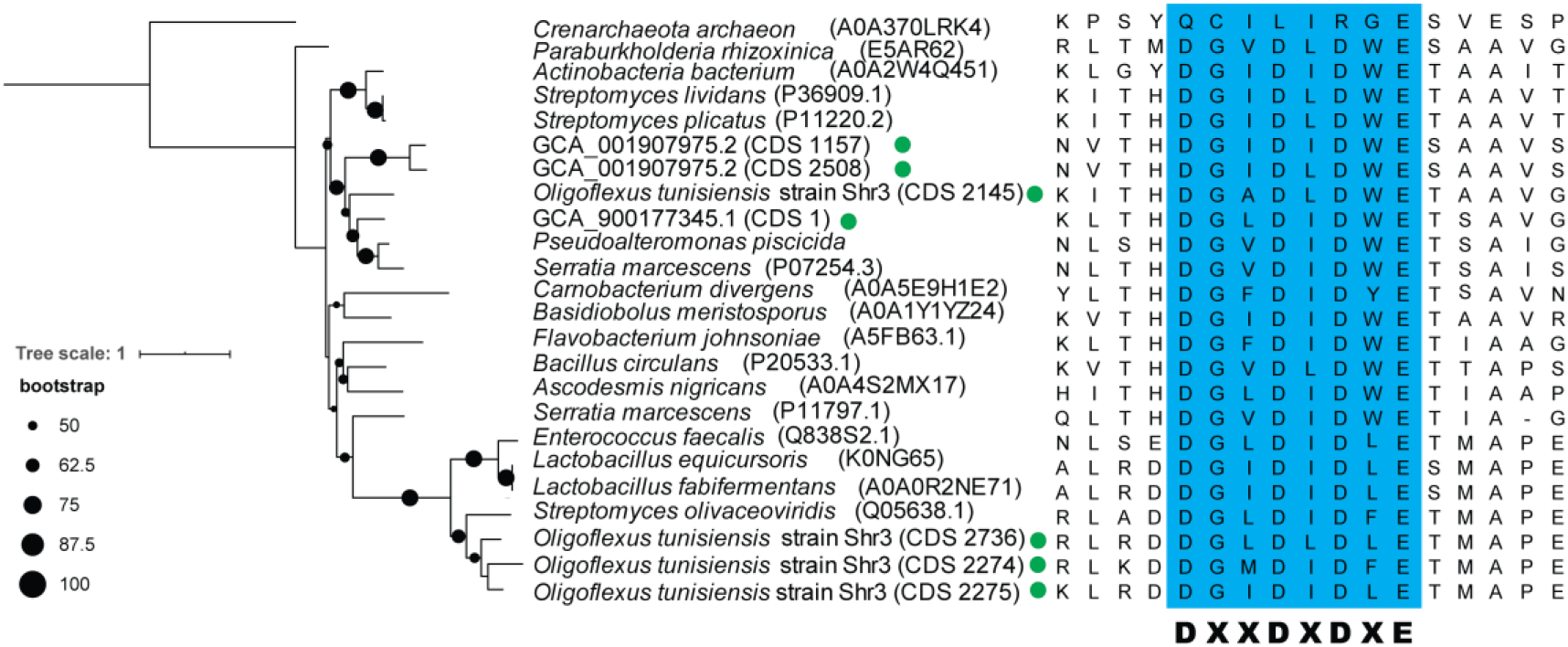
Phylogenetic analysis and conserved motif of chitinases from Oligoflexia. The phylogenetic tree of chitinases was built with RAxML (PROTGAMMALG model). The bootstrap values based on 1,000 replicates were denoted by the size of the black dots on the branches. The Oligoflexia genomes were marked with a green dot. The conserved motif of chitinases was highlighted with blue background in the multiple sequence alignment.

## Conclusion

In this study, Bdellovibrionota was revealed to have four phylogenetic groups, all containing the genes in most of their genomes for two-phase predatory lifestyle. Their vertical distribution in ocean was exhibited. Bacteriovoracia was more prevalent in the deep-sea zones. Genomics traits demonstrated that marine Bdellovibrionota are able to metabolize osmoprotectant and degrade cell wall. The chitinase and chitin-binding protein encoding genes were for the first time focused in Oligoflexia. However, the function and regulation of these genes involved in pathogenic prevention need to be verified by experiments.

## Materials and methods

### Genome collection from public databases

We recruited 148 Bdellovibrionota genomes from GTDB database (26). Eight genomes for model Bdellovibrionota were downloaded from the NCBI. All the genomes were reclassified with GTDB-TK (v.1.0.1). Quality control of all the genomes was conducted using CheckM (v.1.1.2) (67). Mediate-quality genomes used for further analysis were selected with the following standards: 1) Completeness score was more than 70%; 2) Contamination rate was less than 10%; 3) The number of conserved proteins derived from CheckM program was at least 22. Average Nucleotide Identity (ANI) analysis was executed by the pyani (v.0.2.09) (software available from http://widdowquinn.github.io/pyani/) to cluster the genomes with ≽ 98.5% identity (68). The sources of genomes and metagenomes were obtained from NCBI Biosample and Bioproject.

### Samples collection, DNA extraction and sequencing

About 20L marine water samples were *in situ* filtered from the Mariana Trench during *R/V* TS09 by ISMIFF apparatus and sampling procedure had been described in our previous report (69). 20L water samples were also collected at different depths by Niskin bottles from the South China Sea during *R/V* TAN KAH KEE cruise and *R/V* Haigong623 equipped with remotely operated vehicle (ROV) Haixing6000 (Table 1). 0.22μm polycarbonate membranes (Merck Millipore, Bedford, MA) were used for microbial filtration and were then frozen at −80°C degree immediately until use. The polycarbonate membranes were cut into small pieces for DNA extraction using MO BIO Power Soil DNA Isolation Kit (MoBio, Carlsbad, CA, USA) according to the manufacturer’s instruction. The quality and quantity of the DNA extraction were evaluated by 1% agarose gel electrophoresis and Qubit 3.10 Fluorometer (Invitrogen, Life, USA). The good-quality DNA was first sheared randomly to fragments of around 500bp by Focused-ultrasonicattor (Covaris M220) and used for metagenomic library preparation with TruSeq® Nano DNA LT Sample Prep Kit (Illumina, San Diego, CA, USA). The high-throughput sequencing was performed on an Illumina Miseq platform (2 × 300bp).

### Raw data processing, de novo assembly and metagenomes binning

The quality of metagenome raw data was assessed by FastQC (v0.11.8) (70). After Fastp (v0.20.0) (71) treatment, repeated sequences produced by sequencing platform were removed by Fastuniq (72) and then the clean reads were assembled with MEGAHIT (v1.2.5-beta) (73). The contigs >2,000 bp were selected for genome binning with MetaWRAP (v1.2) (models including metabat2, maxbin2 and concoct) (74). The MAGs were then taxonomically classified by gtdb-tk classify (8) and Bdellovibrionota MAGs were selected by the cut-off described above (>70% completeness and <10% contamination).

### Calculation of Bdellovibrionota relative abundance in deep sea

16S rRNA sequences of Bdellovibrionota genomes and MAGs were extracted by using rRNA_hmm_fs_wst (v.0) (75) and used to create a 16S rRNA database. The Bdellovibrionota 16S miTags in Tara Ocean (23) and Mariana (24, 69) marine water metagenomes (Table S7) were identified by blastn (v.2.9.0) (-evalue 1e-05) against the 16S rRNA sequence database of Bdellovibrionota. Only the 16S miTags longer than 100bp and >97% identity to a reference in the database were selected for further calculation of their relative abundance in the marine samples. The Tara Ocean data were downloaded from http://ocean-microbiome.embl.de/companion.html. T-test was performed for significance analysis.

### Genome annotation and metabolic reconstruction

The high-quality genomes from public databases (>90% completeness and <5% contaminant) and our five MAGs were used for gene annotation. ORFs were predicted by Prodigal (v.2.6.3) (76) and were then annotated by Kofamscan (77) (v. 1.0.0; -f mapper) against KEGG database (release 92) and by BLASTP (v.2.9.0) against COG database (COG_2019_v11.0). Carbohydrate related enzymes were searched with hmmscan (v.3.2.1) against CAZy database (dbCAN-HMMdb-V7). Peptidases were identified by BLASTP against MEROPS database (28) (pepunit.lib; -id 50; -e 1e-10). Wilcoxon test was conducted by R package to identify the genes significantly different in abundance between different groups.

### Phylogenetic analysis of conserved proteins and 16S rRNA genes

The marker genes of genomes were aligned using hmmerAlign.py (called by CheckM). The alignment of 43 concatenated conserved proteins was used for reconstruction of phylogenomic tree with RAxML (78) (v.8.2.12; -# 1000; -m PROTGAMMALG) after optimization with trimAl (79) (v.1.4). All the 16S rRNA sequences extracted by rRNA_hmm (75) were treated with CD-HIT (80) (v.4.8.1; -c 0.97; -n 10; -G 1; -M 10000) to remove the same species. Two 16S rRNA sequences of *E. coli* were added as an out-group. All 16S rRNA sequences were aligned with MAFFT (81) (v.7.407) and optimized with trimAl. RAxML (v.8.2.12; GTRCAT model) was used to build a 16S rRNA phylogenetic tree with bootstrap values based on 1000 replicates. All phylogenetic trees were visualized with iTOL (https://itol.embl.de/).

## Acknowledgements

We are grateful to the team members aboard the *R/V* TS09 for their invaluable efforts in the sampling cruise. Great thanks are given to J. Li, J. Chen, S.X. Wang, Y.Z. Xin and D.S. Cai for their ISMIFF sampling. This study was supported by Key Research and Development Program of Hainan Province (E050010406) and the National Key Research and Development Program of China (2018YFC0310005 and 2016YFC0302501).

---

**Figure S1.**
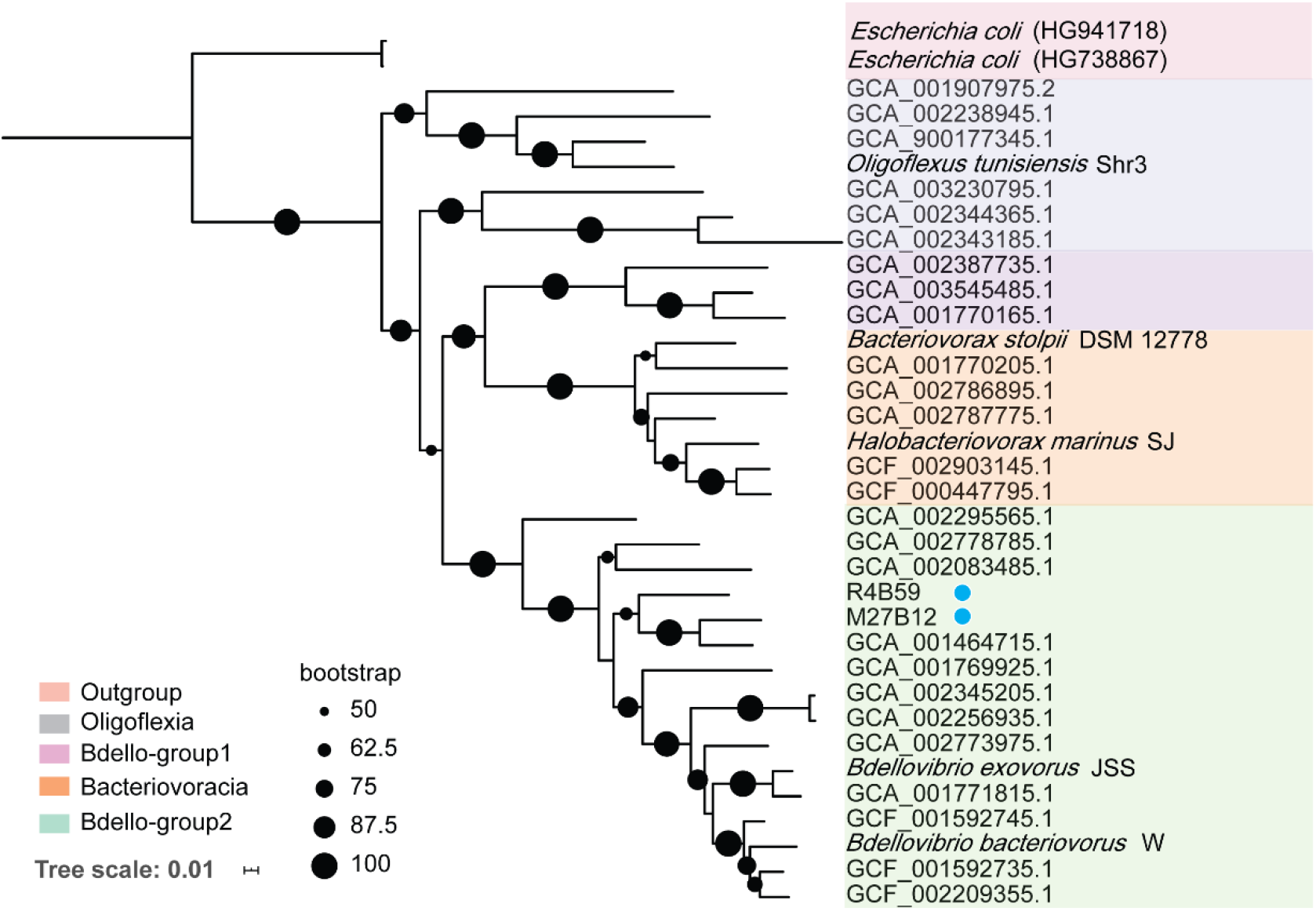
Rooted phylogenetic tree of Bdellovibrionota 16S rRNA genes. Maximum-likelihood (ML) phylogenetic tree of 16S rRNA genes extracted from high-quality genomes of Bdellovibrionota were reconstructed using PROTGAMMALG model. Blue dots refer to the MAGs retrieved from marine waters of this study.

**Figure S2.**
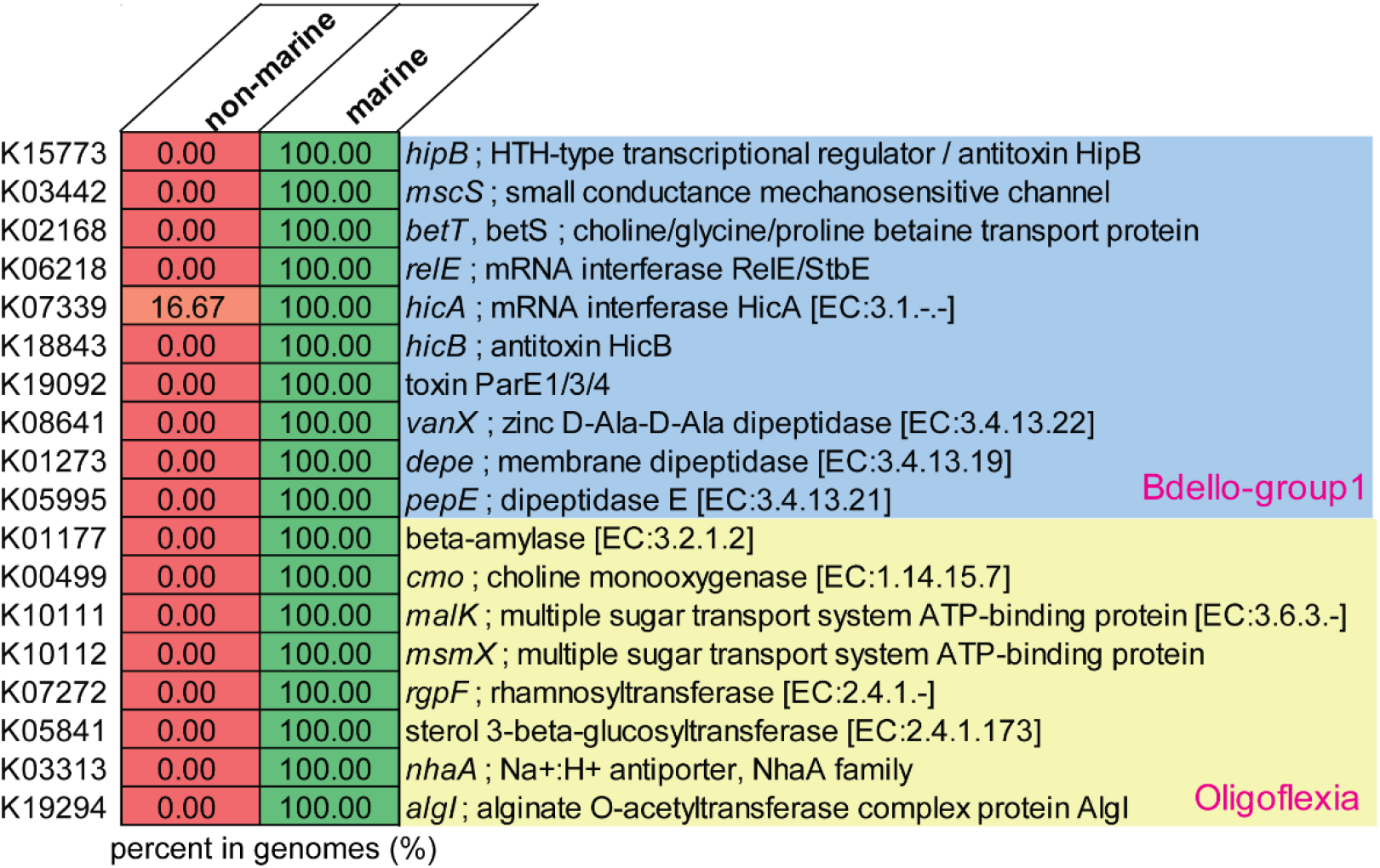
Genes in genomes affiliated with Oligoflexia and Bdello-group1 from marine environment. The percentage of the KEGG genes in the genomes was calculated for the marine and non-marine genomes of Oligoflexia and Bdello-group1. Genes identified in Bdello-group1 marine genomes were involved in betaine metabolism, type II toxin-antitoxin (TA) system and dipeptidases. Genes present in Oligoflexia marine genomes were involved in cold resistance (beta-amylase), betaine synthesis (*cmo*), multiple sugar transport and metabolism (*malK, msmK, algI* and *rgpF*).

**Figure S3.**
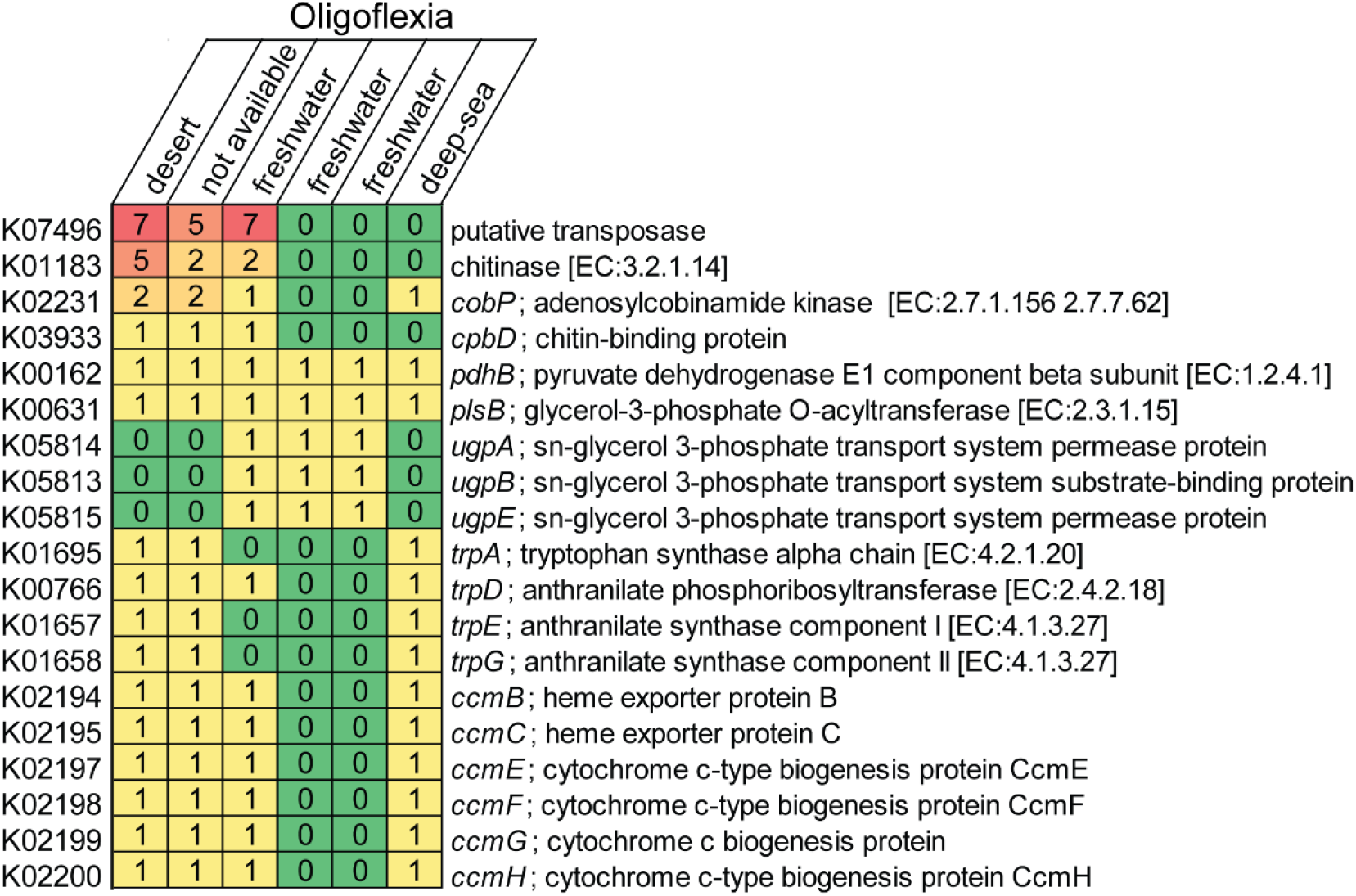
Copy number of interested KEGG genes of Oligoflexia from different environments. The special genes are only present in Oligoflexia genomes inhabiting different environments. Chitinase (more than one copy) and chitin-binding protein coding genes were predicted in some genomes.

**Figure S4.**
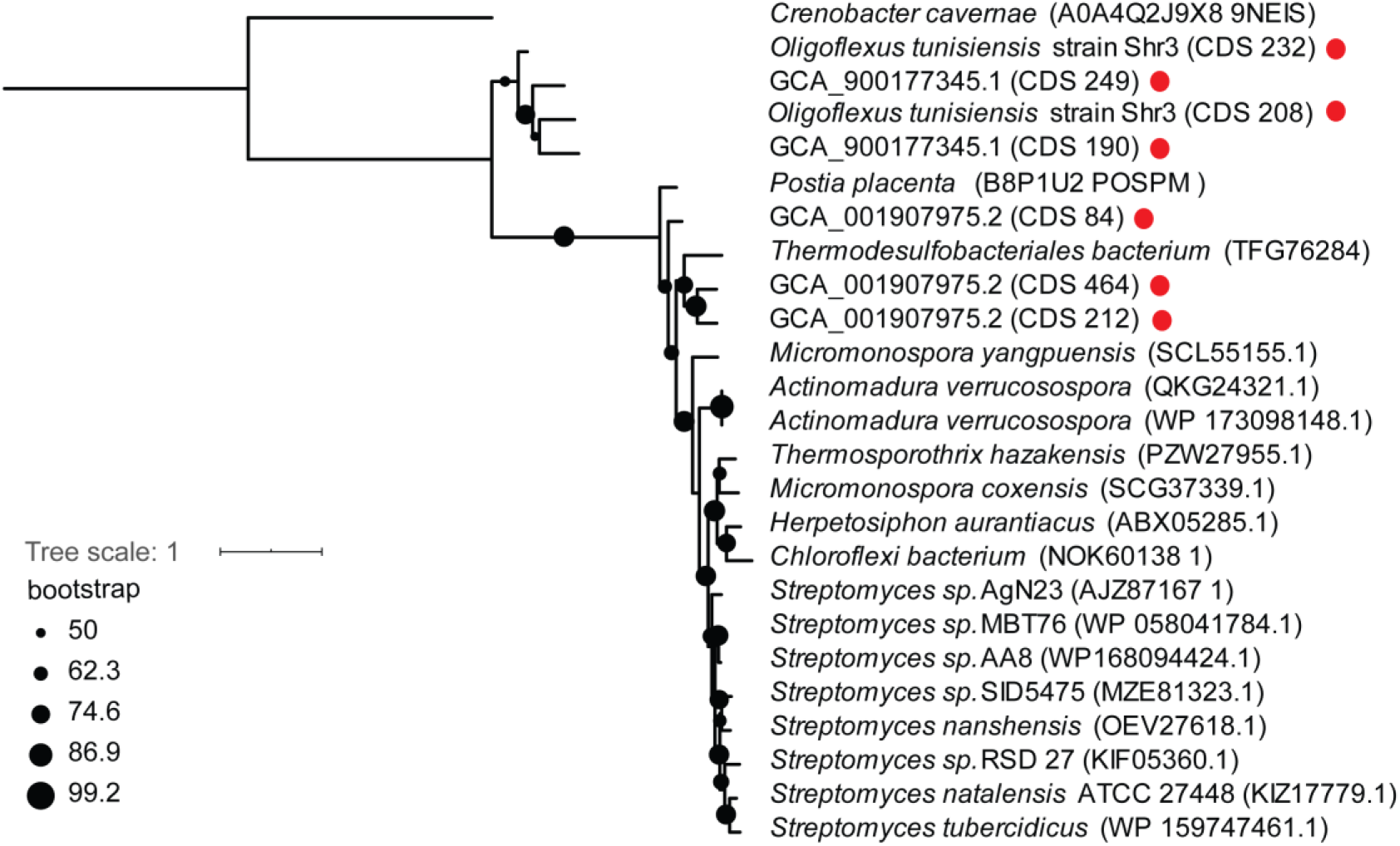
Phylogenetic tree of chitin-binding proteins encoded by Oligoflexia genomes. The red dots refer to amino acid sequences of chitin-binding proteins predicted based on KEGG annotation of Oligoflexia genomes.

Table S1. GTDB annotation of all Bdellovibrionota genomes from GTDB-TK database, five MAGs of our marine water samples and eight Bdellovibrionota genomes from NCBI.

Table S2. Information of completeness, contamination, genome size and GC content of all genomes.

Table S3. Average nucleotide identity of all genomes.

Table S4. Environment source of genomes.

Table S5. KEGG and COG genes of Bdellovibrionota involved as predator.

Table S6. KEGG annotation results of genomes.

Table S7. The information of water samples collected from Mariana Trench with the depth no less than 1,000m.

Table S8. Blastp results of chitinase and chitin-binding protein.

